# Widespread signatures of negative selection in the genetic architecture of human complex traits

**DOI:** 10.1101/145755

**Authors:** Jian Zeng, Ronald de Vlaming, Yang Wu, Matthew R Robinson, Luke Lloyd-Jones, Loic Yengo, Chloe Yap, Angli Xue, Julia Sidorenko, Allan F McRae, Joseph E Powell, Grant W Montgomery, Andres Metspalu, Tonu Esko, Greg Gibson, Naomi R Wray, Peter M Visscher, Jian Yang

**Affiliations:** Institute for Molecular Bioscience, The University of Queensland, Brisbane, Queensland 4072, Australia; Department of Complex Trait Genetics, VU University, Center for Neurogenomics and Cognitive Research, Amsterdam, 1081 HV, The Netherlands; Erasmus University Rotterdam Institute for Behavior and Biology, Rotterdam, 3062 PA, The Netherlands; Department of Computational Biology, University of Lausanne, 1011 Lausanne, Switzerland; Estonian Genome Center, University of Tartu, Tartu, Estonia; School of Biology and Center for Integrative Genomics, Georgia Institute of Technology, Atlanta, GA 30332, USA; Queensland Brain Institute, The University of Queensland, Brisbane, Queensland 4072, Australia

## Abstract

Estimation of the joint distribution of effect size and minor allele frequency (MAF) for genetic variants is important for understanding the genetic basis of complex trait variation and can be used to detect signature of natural selection. We develop a Bayesian mixed linear model that simultaneously estimates SNP-based heritability, polygenicity (i.e. the proportion of SNPs with nonzero effects) and the relationship between effect size and MAF for complex traits in conventionally unrelated individuals using genome-wide SNP data. We apply the method to 28 complex traits in the UK Biobank data (*N* = 126,752), and show that on average across 28 traits, 6% of SNPs have nonzero effects, which in total explain 22% of phenotypic variance. We detect significant (p < 0.05/28 =1.8×10^−3^) signatures of natural selection for 23 out of 28 traits including reproductive, cardiovascular, and anthropometric traits, as well as educational attainment. We further apply the method to 27,869 gene expression traits (*N* = 1,748), and identify 30 genes that show significant (p < 2.3×10^−6^) evidence of natural selection. All the significant estimates of the relationship between effect size and MAF in either complex traits or gene expression traits are consistent with a model of negative selection, as confirmed by forward simulation. We conclude that natural selection acts pervasively on human complex traits shaping genetic variation in the form of negative selection.

## Introduction

Dissecting the genetic architecture of complex traits is important for understanding the genetic basis of phenotypic variation and evolution. For a complex trait that influences fitness, natural selection plays an important role in shaping its genetic architecture^1^, which in turn provides information to infer the action of natural selection. Given most traits are polygenic, natural selection is likely to act simultaneously on many trait-associated variants that have pleiotropic effects on fitness (known as polygenic selection^2–4^). Unlike a selective sweep model^5^ where there are often a limited number of mutations under relatively strong selection, it is difficult to detect the signals of polygenic selection due to the selection pressure being spread over many variants of small effect. However, evidence for natural selection can be inferred from the relationship between effect size and minor allele frequency (MAF) at the genome-wide variants. For example, mutations that are deleterious to fitness are selected against and thus kept at low frequencies by negative selection, resulting in a correlation between effect sizes and MAF^6,7^. The estimation of the joint distribution of effect size and MAF can be used to detect signature of natural selection and thereby to infer the relationship between a complex trait and fitness.

Genome-wide association studies (GWAS) have detected thousands of SNPs associated with complex traits, which have helped to characterize the genetic architecture of these traits^8–13^. However, the genome-wide significant SNPs discovered in GWAS jointly tend to explain only a fraction of the heritability as many SNPs with small effects yet to be detected^14^. Furthermore, a proportion is missed due to the incomplete linkage disequilibrium (LD) between causal variants and SNP markers^14^. To address the “missing heritability” problem^14,15^ in GWAS, mixed linear model (MLM) approaches have been used to estimate the genetic variance explained by all SNPs used in a GWAS. GREML is a prevailing class of MLM-based approaches where all SNP effects are fitted together as random effects^16^. GREML analyses using common SNPs (MAF > 0.01) have uncovered a large proportion of the “missing heritability” for height^17^, BMI^17^ and psychiatric disorders^18^. The GREML method assumes that all SNPs have an effect on the trait^16^ and thus does not allow us to estimate the degree of polygenicity (i.e. the proportion of SNPs with nonzero effects). Bayesian multiple regression is another class of MLM-based methods that enable us to make posterior inference about polygenicity by assuming SNP effects are drawn from a mixture distribution of zero and nonzero components^19–21^. Bayesian MLM methods have been widely used in livestock and plant breeding^22^ and have attracted increasing attention in humans for characterizing the genetic architecture of complex traits and diseases^20,23,24^. However, neither GREML nor Bayesian MLM approaches explicitly model the relationship between effect size and MAF, an important characteristic of the genetic architecture for complex traits. This relationship can be used to detect signatures of natural selection^7,25^ and inform the design of future genetic mapping studies.

In this study, we developed an MLM-based Bayesian method that can simultaneously estimate SNP-based heritability, polygenicity and the joint distribution of effect size and MAF in conventionally unrelated individuals using GWAS data. We applied the method to 28 complex traits in the UK Biobank (UKB) data^26^, and 27,869 gene expression traits in the Consortium for the Architecture of Gene Expression (CAGE) dataset^27^, and identified a number of complex traits and gene expression traits for which there is significant evidence of natural selection on the associated SNPs.

## Results

### Method overview

Under the Bayesian MLM framework, we propose to model the relationship between effect size and MAF using the following mixture distribution as prior for each SNP effect

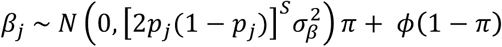

where *β*_*j*_ is the allelic substitution effect of a SNP *j, p*_*j*_ is the MAF of the SNP, 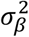 is a constant factor (i.e. variance of SNP effects under a neutral model), *ϕ* is a point mass at zero, and *π* is the proportion of SNPs with nonzero effects (polygenicity). The variance of the effect size of SNP *j* is 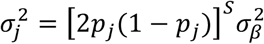., which is a function of MAF of the SNP. Thus, the parameter *S* measures the relationship between effect size and MAF. If *S* = 0, the effect size is independent of MAF (neutral model). If there is selection, the effect size can be positively (*S* > 0) or negatively (*S* < 0) related to MAF. All these parameters are treated as unknown with appropriate priors (**Online Methods**). Our model (referred to as BayesS) allows simultaneous estimation of multiple characteristics of the genetic architecture: SNP-based heritability (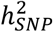) polygenicity (π) and the relationship between SNP effect and MAF (5). We use a gradient-based sampling algorithm, Hamiltonian Monte Carlo^28^, to sample *5* from the posterior distribution, and use Gibbs sampling for other parameters in the model by assuming conjugate priors. Furthermore, we use a parallel computing strategy following Fernando *et al.*^29^ to allow the analysis to be scalable to very large samples sizes *(N >* 100,000). Details of sampling scheme and parallel computing strategies are given in the Supplementary Note.

In the hypothesis test against *S* = 0, we used two approaches to control false positives. The first approach is to control the family-wise type I error rate (FWER) using the theory that the posterior mode standardized by the posterior standard error (s.e.) asymptotically follows a standard normal distribution under the null^30^. The asymptotic normality of the posterior distribution was justified by simulation with the UKB cohort (Supplementary Fig. 1). The second approach is to control the proportion of false positives^31^ (PFP) among rejections (also known as the marginal false discovery rate or mFDR^32^) based on the posterior probability given the data, *e.g.* Pr(*S* < 0**|**𝒟)**Supplementary Note**). We show by simulation that if the true distribution of *S* is used as the prior, then rejecting *S* = 0 with Pr(*S* < 0**|**𝒟) ≥ *γ* guarantees PFP or mFDR to be less than 1 — *γ* (**Supplementary Fig. 2**). The former approach is more stringent but the advantage of the latter approach is that the power is not inversely related to the number of traits (tests)^31^.

### Assessing the robustness of parameter estimation through simulations based on real genotype data

We used simulations based on real GWAS genotype data from the Atherosclerosis Risk in Communities (ARIC) study^33^ to assess our method in estimating the parameters 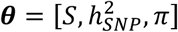. The ARIC data consist of 12,942 unrelated individuals and 564,959 Affymetrix SNPs with MAF > 1% after quality control (**Online Methods**). In our simulation, 1,000 SNPs were chosen at random to be causal variants, with their effects related to MAF through an *S* value ranging from -1 to 1 in different scenarios (**Online Methods**). Since the number of causal variants was known, polygenicity was assessed by the number of SNPs with nonzero effects *(m*_*NZ*_). Based on the Markov chain Monte Carlo (MCMC) samples, the point estimate (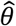), standard error (s.e.) or credible interval for each parameter was given by the mode, standard deviation (s.d.) or highest probability density (HPD) of its posterior distribution, respectively.

Results (**Fig. 1**) show that when both causal variants and SNP markers were fitted in the analysis, *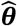* from BayesS was unbiased with respect to the true parameters. When the causal variants were not included in the analysis, both 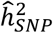 and the absolute value of *Ŝ* were slightly underestimated, due to imperfect tagging, a similar issue as discussed in Yang *et al.*^16^. For polygenicity, however, *m*_*NZ*_ estimate tended to be larger than the number of causal variants, probably because some causal variants could be better tagged by multiple SNPs. Thus, in practice, *π* should be interpreted as the proportion of non-null SNPs, which is likely to be larger than the proportion of causal variants. Results also show that the s.e. for *Ŝ*, 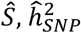 and 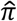 is consistent with the s.d. of the estimates from 100 simulation replicates (**Supplementary Table 1**).

**Figure 1:**
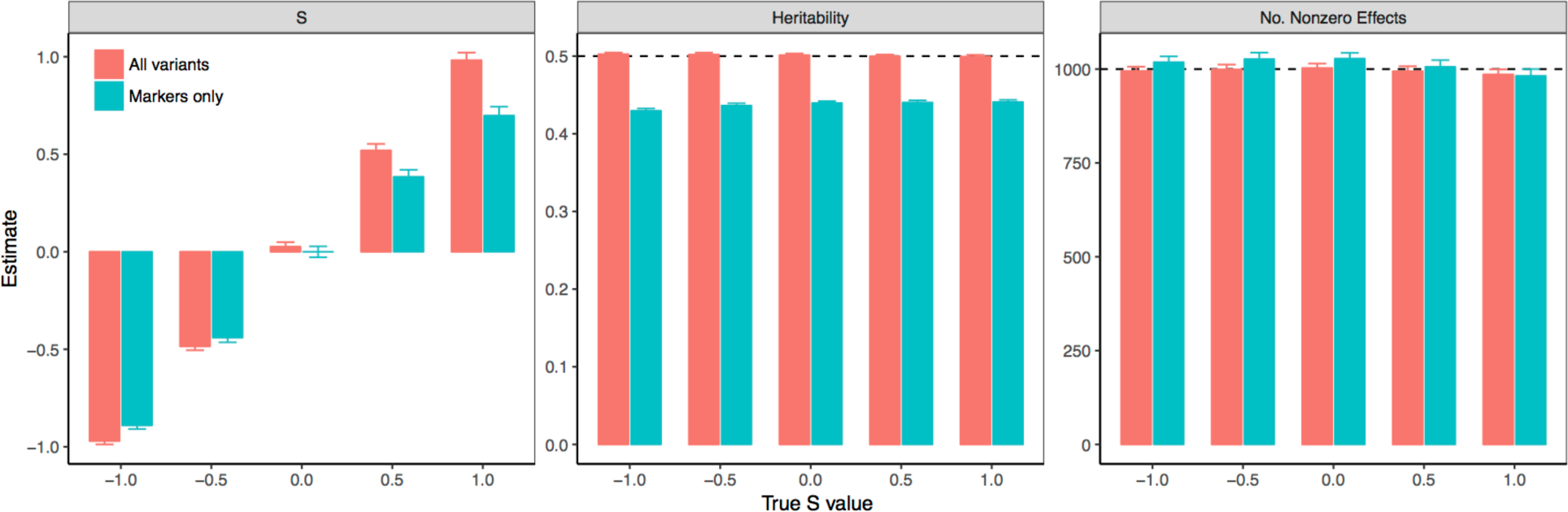
Estimation of the genetic architecture parameters, e.g. *S*, heritability and polygenicity, for a simulated trait using the ARIC data. Results are the mean estimates with s.e.m. (cap) over 100 simulation replicates for a spectrum of *S* parameter values. Colour indicates the results of BayesS with both causal variants and SNP markers (red) or with SNP markers only (blue). The heritability at the 1,000 randomly selected causal variants was 0.5. The number of nonzero effects is the number of SNPs with nonzero effects averaged over MCMC iterations.

### Analysis of 28 complex traits in the UK Biobank data

We applied the BayesS method to 36 complex traits on 126,545 unrelated individuals of European ancestry in the UKB^26^ with 483,634 Affymetrix SNPs (MAF > 1%) after quality controls (**Online Methods**). Out of the 36 traits 21 have *N >* 100,000. Two commonly used long-chain diagnostic tests were adopted to assess the convergence of the MCMC algorithm (**Supplementary Note**). Traits with results that did not pass our convergence tests were those with the smallest sample sizes, 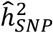 close to zero, or both (**Supplementary Fig. 3**). We focus on the results of 28 traits that passed both convergence tests for all of the three genetic architecture parameters. These traits include 24 quantitative traits, 2 diseases: major depressive disorder (MDD) and type 2 diabetes (T2D), and 2 categorical traits: male pattern baldness (MPB) and years of schooling (educational attainment). **Supplementary Fig. 4** shows the distributions of the estimates across these traits for the three genetic architecture parameters.

### Comparison of the genetic architecture between Height and BMI

The genetic architectures of height and BMI have been relatively well studied compared to other complex traits^34–40^. Thus, it is interesting to compare our results for height and BMI **(Fig. 2)** with the previous findings. Both traits have a large sample size in the UKB: *N* = 126,545 for height and *N* = 126,389 for BMI. For both traits, a negative *S* was detected with extremely high significance level(*P* =1.8 × 10^−106^ for height and *P* =2.8 × 10^−14^for BMI), meaning that lower-MAF variants tend to have larger effect size (absolute values). These results suggest that both height- and BMI-associated SNPs have been under selection, in line with the conclusions drawn from two recent studies^36,39^. The posterior mode of *S* was -0.422 (s.e. = 0.019) for height, remarkably lower than that of -0.295 (s.e. = 0.039) for BMI, suggesting that the proportion of genetic variation attributable to SNPs with low MAF for height is larger than that for BMI, consistent with the result from a previous study^36^. These results also imply that overall height-associated SNPs are under stronger selection than BMI-associated SNPs. The phenotypic variance explained by common SNPs (MAF > 0.01) was 52.8% (s.e. = 0.3%) for height and 27.7% (s.e. = 0.4%) for BMI, consistent with the estimates of 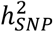 for height and BMI based on common SNPs reported previously^34–36^. The posterior distribution of *π* provides an estimate of 4.8% (s.e. = 0.1%) of SNPs having nonzero effects on height, significantly lower than that of 9.4% (s.e. = 0.5%) for BMI. These results suggest that BMI is more polygenic but less heritable than height, consistent with the results from a recent study using BayesR, a Bayesian multicomponent mixture model^41^. As a consequence, a BMI-associated SNP on average would explain a relatively smaller proportion of 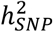, compared with a height-associated SNP, which may explain the higher uncertainty in the estimates of the hyperparameters such as *S* and *π* for BMI. These results also explain why the number of genome-wide significant SNPs *(m*_*GWS*_) identified from the GIANT meta-analysis for BMI (*m*_*GWS*_ *=* 97) was smaller than that for height (*m*_*GWS*_ *=* 697) despite the fact that the sample size for BMI (N = _~_ 340,000)^35^ was considerably larger than that for height (N = _~_250,000)^34^.

**Figure 2:**
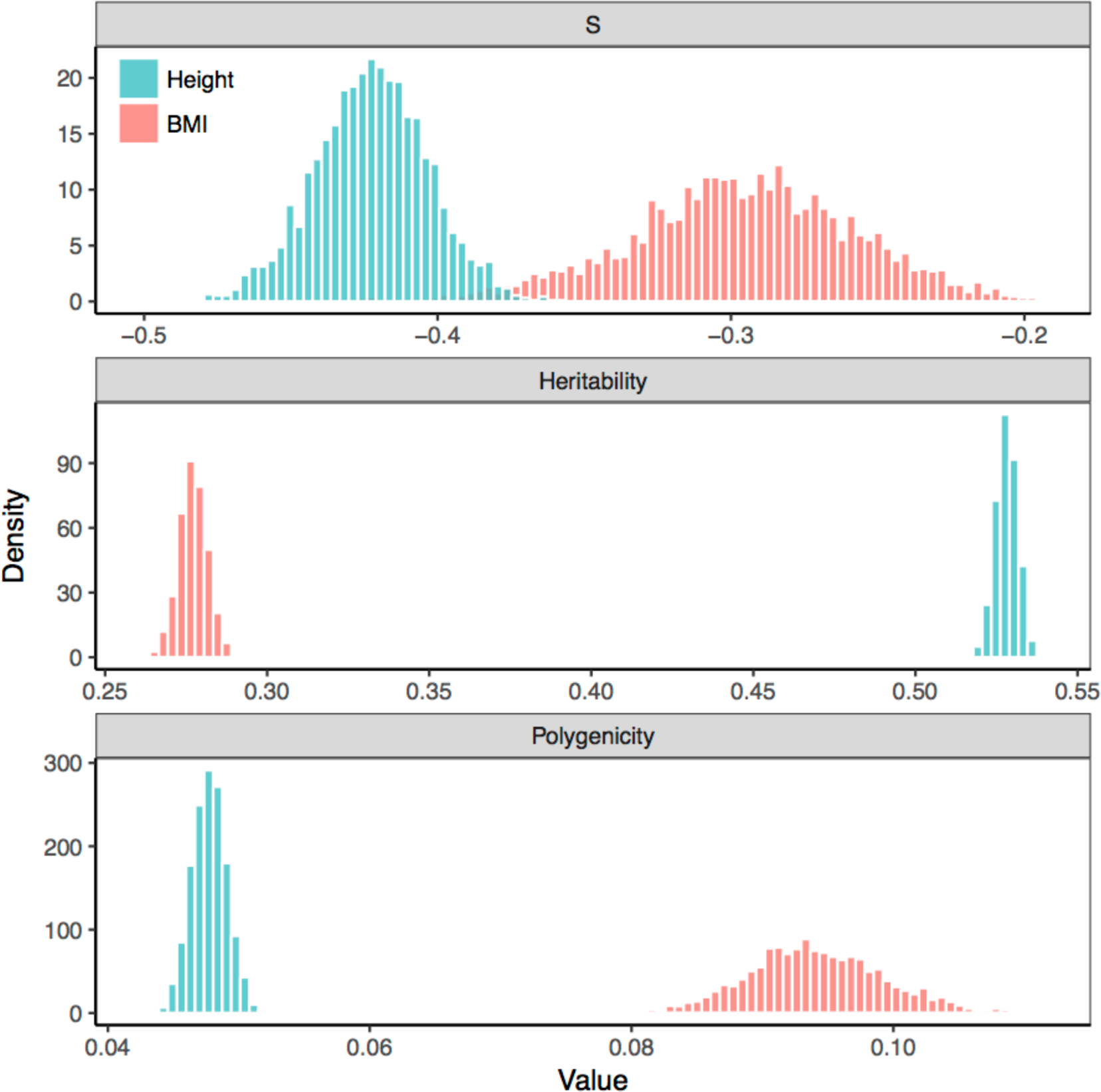
Posterior distributions of the genetic architecture parameters for height versus BMI using data from UKB. S measures the relationship between SNP effect size and MAF. Polygenicity is defined as the proportion of SNPs with nonzero effects.

### Inference on natural selection

Of the 28 traits that passed our convergence tests, 23 traits (including reproductive, cardiovascular and anthropometric traits and educational attainment) had significant negative *S* estimates with Pr (*S* < 0**|**𝒟) = 1 and *P* < 0.05/28 (**Supplementary Table 2**), providing strong evidence that these traits have been under selection. The estimates of *5* over traits ranged from -0.601 (age at menopause) to 0.016 (fluid intelligence score) with mean -0.348, median -0.364 and s.d. 0.112. Interestingly, all the significant estimates of *S* were negative (see below for forward simulation to infer the type of selection from the sign of *S*). The magnitudes of *Ŝ*, *i.e.* |*Ŝ*|, reflects the strength of selection on the trait-associated SNPs. Traits with the largest |*Ŝ*| are related to fertility and heart function (**Fig. 3**), including age at menopause (*Ŝ* = −0.601, s.e. = 0.073), pulse rate (*Ŝ* = −0.481, s.e. = 0.048), waist circumference adjusted for BMI (WCadjBMI, *Ŝ* = −0.436, s.e. = 0.036) and waist-hip ratio adjusted for BMI (WHRadjBMI, *Ŝ* = −0.436, s.e. = 0.049). It has been reported that WCadjBMI and WHRadjBMI are associated with cardiovascular events^42^, and WHRadjBMI is strongly correlated with pregnancy rate^43^. Other reproductive and cardiovascular traits, such as age at first live birth, age at menarche and blood pressure, had relatively high |*Ŝ*| as well. Thus, our results suggest that reproductive and cardiovascular traits are closely related to fitness and the SNPs that are associated with these traits have been under relatively stronger selection than SNPs associated with other traits.

**Figure 3:**
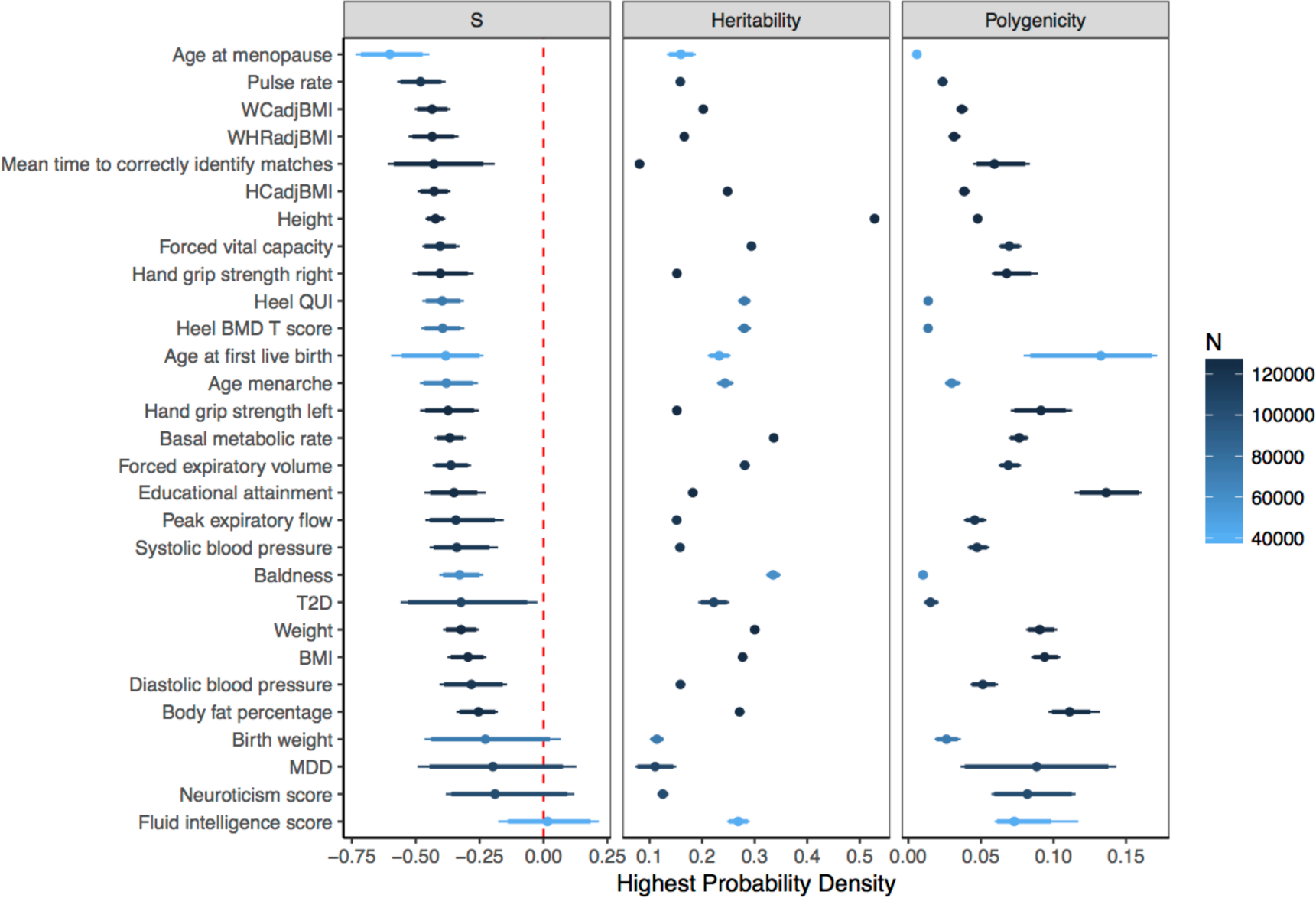
Posterior modes with credible intervals for the genetic architecture parameters using BayesS. Results are for the 28 UKB complex traits that had passed convergence tests on the MCMC chain. The bold line represents 95% credible interval (highest posterior density, HPD) and the thin line represents 90% credible interval. Sample size *N* for each trait is shown by the colour gradient. Polygenicity is defined as the proportion of genome-wide SNPs with nonzero effects on the trait.

Height (*Ŝ* = −0.422), handgrip strength (right: -0.404, left: -0.374), lung function related traits (-0.405 – -0.362), heel bone mineral density (-0.394) and basal metabolic rate (-0.367) had a moderate to high |*Ŝ*| (**Fig. 3** and **Supplementary Table 2).** Evidence of selection for height has been reported from multiple studies using different approaches^36–40^. The two diseases, MDD and T2D, had negative *Ŝ* but the P-values did not reach FWER significance threshold, although the posterior probability of *S* < 0 for T2D was as high as 0.983. However, the power to detect a significant *Ŝ* may not be comparable to those quantitative traits, given the number of cases was less than 10,000 for each. A recent large-scale GWAS based on whole-genome sequencing data also did not detect a signal of selection on T2D-associated variants^8^. Fluid intelligence score is the only trait with *Ŝ* at almost zero (*Ŝ* = 0.016, s.e. = 0.096), which seems to suggest that fluid intelligence (FI) is not pertinent to fitness. However, there is strong evidence of negative selection on the SNPs associated with educational attainment (EA, *Ŝ* = −0.350, s.e. = 0.055), which is thought to be a proxy of intelligence. Indeed, the genetic correlation between EA and FI was as high as 0.665 (s.e. = 0.052) estimated from a bivariate LD score regression^44^. Thus, it may be due to the limited statistical power that we did not detect the signal of selection for FI.

For traits with a significant estimate of *S*, we demonstrated the relationship between effect size and MAF by a plot of the cumulative genetic variance explained by SNPs (*V*_g_) against MAF (**Fig. 4**), where MCMC samples of SNP effects were used to compute *V*_g_ for SNPs with MAF smaller than a threshold on the x-axis (**Supplementary Note**). Under an evolutionarily neutral model, *V*_g_ is linearly proportional to MAF^45^ (diagonal line), therefore the area under the curve (AUC) is 0.5. All traits with significant estimates of had the curve of cumulative genetic variance above the diagonal line, with |*Ŝ*| highly correlated with the AUC *(r* = 0.902), an alternative way of illustrating the evidence of natural selection.

**Figure 4:**
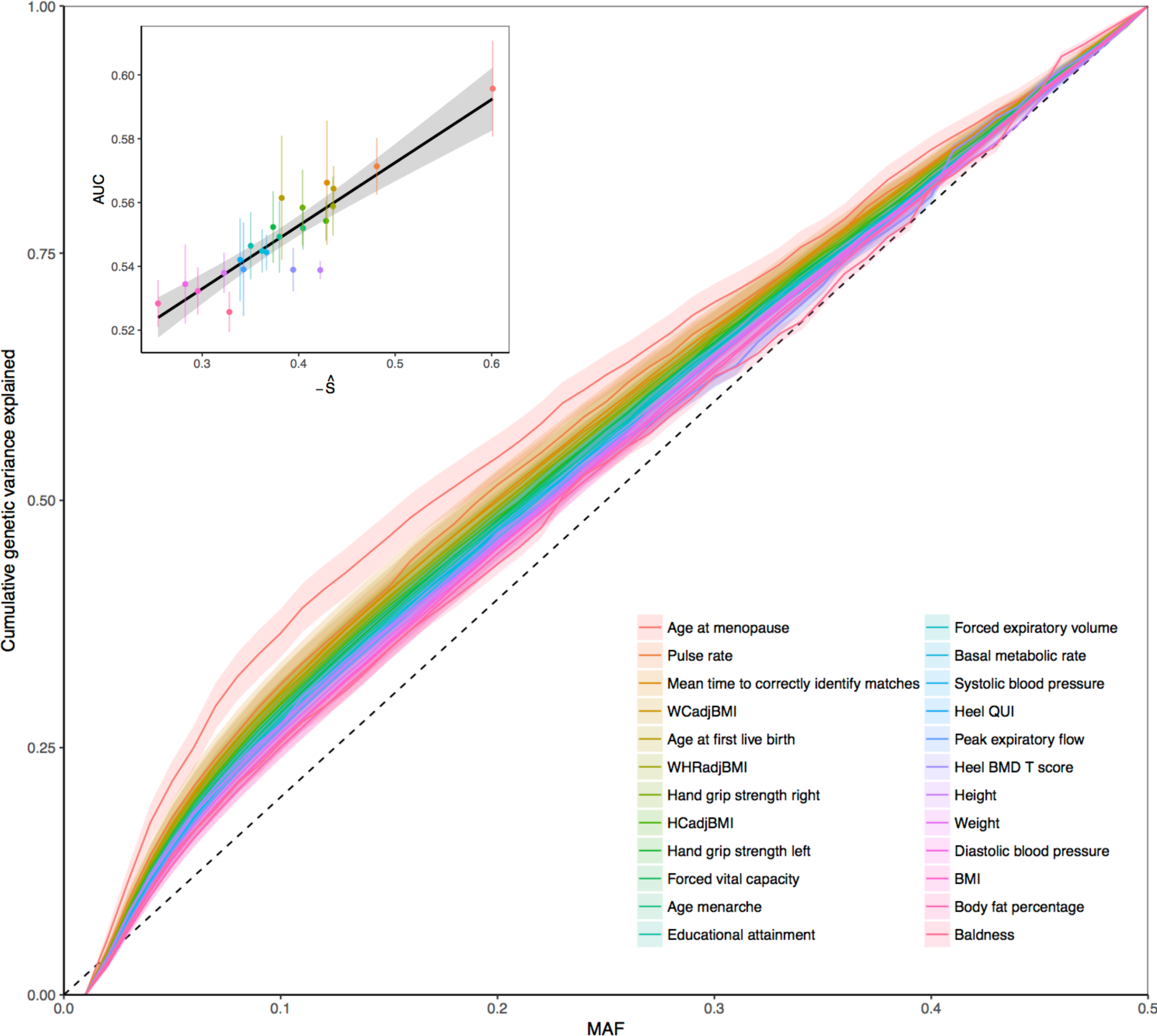
Cumulative genetic variance explained by SNPs with MAF smaller than a threshold on the *x*-axis. The lines are the posterior means for the 23 UKB complex traits from UKB for which the estimates of *S* were significantly different from zero. Shadow shows the posterior standard error. The inner graph shows the relationship between the area under the curve (AUC) of the cumulative genetic variance and negative *Ŝ* (bar shows s.e.) across traits.

### Inference on SNP-based heritability

The 28 traits had low to moderate estimates of *h*^2^_SNP_ with mean 0.221, median 0.212, and s.d. 0.093, and were all significantly above zero (**Supplementary Table 3**). Note that traits with 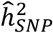 close to zero had failed in MCMC convergence tests, therefore the mean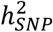 estimate across traits is likely to be inflated. For MDD (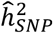 = 0.111, s.e. = 0.021) and T2D (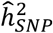 = 0.222, s.e. = 0.015), the estimates were on the liability scale and were converted from the observed scale ^15^, assuming a population prevalence of 15%^46^ and 3%^47^, respectively. The sorted estimates across traits are shown in **Supplementary Fig. 5**. Besides height (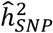 = 0.528), traits with the highest 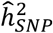 include basal metabolic rate (0.336), which has been reported to be 0.2–0.4 in model animals^48^, and MPB (0.335), which has been reported to be a highly heritable trait in both pedigree^49^ and genomic^50^ analysis. Traits with the lowest 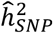 include mean time to correctly identify matches (0.081), MDD (0.111), birth weight (0.114) and neuroticism score (0.125), in line with the low estimates of h._NP_ from previous studies in MDD^51^ and neuroticism score^52^. Given that most published estimates were obtained using whole-genome imputed SNPs, they are likely to be slightly higher than our estimates that are only based on the SNPs on Affymetrix Axiom Genotyping Arrays. For example, a recent study^53^ on educational attainment in UKB gave an estimate of 0.21 (s.e. = 0.006), slightly higher than our estimate of 0.182 (s.e. = 0.004). Our estimate of 0.528 (s.e. = 0.003) for height is slightly but not significantly lower than that of 0.56 (s.e. = 0.023) in Yang *etal.*^36^. For BMI, our estimate of 0.277 (s.e. = 0.004) is highly consistent with that of 27% (s.e. = 2.5%) in Yang *et al.*^36^. Across traits, 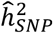 seems to be independent of either *Ŝ or* 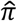 but the s.e. of *Ŝ* and 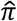 decrease as 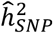 increases (**Supplementary Fig. 6**).

### Inference on polygenicity

The distribution of 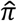 had mean 5.9%, median 5.5% and s.d. 3.6% across traits, and ranged from 0.6% (s.e. = 0.1%) to 13.6% (s.e. = 1.3%) (**Supplementary Table 4**). This suggests that all the 28 complex traits are polygenic with ~30,000 common SNPs with nonzero effects on average. Note that our simulation above suggests that this is likely to be an overestimation of the number of causal variants (**Fig. 1**). Interestingly, age at menopause, the trait with highest magnitude of *S* (-0.601), had the lowest estimate of polygenicity 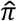 (0.6%, s.e. = 0.1%) (**Supplementary Fig. 5**). Educational attainment had the highest 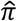 (13.6%, s.e. = 1.3%), which is reasonable because it is a compound trait of several sub-phenotypes so that many SNPs have an effect. It is followed by age at first live birth (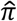*=* 13.3%, s.e. = 2.5%), body fat percentage (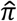*=* 11.1%, s.e. = 0.8%) and BMI (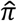*=* 9.4%, s.e. = 0.5%). On the contrary, these traits had low to moderate magnitude of *Ŝ.*

### Analysis of gene expression traits in the CAGE data

Analysing expression levels of all probes in the CAGE dataset^27^ (1,748 unrelated individuals of European ancestry) using the standard BayesS approach would be computationally challenging, as it would require us to perform 36,778 distinct BayesS analyses. However, given that many probes have a very limited number of associated SNPs, we developed a nested version of the BayesS model. This nested approach speeds up the analyses by considering SNPs in proximity collectively as a window, which allows for fast “jumping” over windows with zero effect (**Online Methods**). We showed by simulation that the nested model produces similar results as the standard BayesS approach in the analyses of both simulated (**Supplementary Fig. 7**) and UKB data (**Supplementary Fig. 8**) while being six times as fast as the standard BayesS approach for traits with low polygenicity (**Supplementary Fig. 9**). Using the nested BayesS model, we were able to fit 1,066,738 imputed SNPs (MAF > 1% and in common with those on HapMap3^54^) for the gene expression traits by partitioning the genome into 12,937 non-overlapping 200-Kb segments. Thus, the polygenicity (*π*) is interpreted as the proportion of segments with nonzero effects in nested BayesS.

After convergence tests, 27,869 probes remained, most of which had low 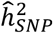 (mean = 0.147, median = 0.122 and s.d. = 0.088) and polygenic architecture (mean *π* = median *π* = 5.2% and s.d. = 2.3%) (**Supplementary Fig. 10**). With unrelated individuals only, our 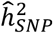 were moderately correlated with the GREML estimates *(r* = 0.568, **Supplementary Fig. 11**) despite the relatively small sample size. The estimates of polygenicity 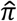 suggest widespread trans-regulatory effects on gene expression in humans. To identify genes under selection, we mapped 21,303 out of the 27,869 probes to the genome with at least “good” probe annotation quality^55^, which tagged 15,615 genes. Applying a Bonferroni correction for the number of probes mapped to the genome, we identified 32 probes that had *Ŝ* significantly different from zero (*P* < 0.05/ 21,303 = 2.3×10^−6^; **Fig. 5** and **Supplementary Table 5**). These probes were mapped to 30 unique genes (Fig. 6) and all had negative *Ŝ* (mean = -1.259, s.d. = 0.185), moderate 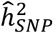 (mean = 0.412, s.d. = 0.075) and small 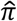 (mean = 0.0268, s.d. = 0.011). The alternative approach to control false positives is to limit PFP, which is less stringent but more powerful compared with limiting FWER. With this approach, a number of additional probes were identified with Pr(*S* < 0|𝒟) ≥ 0.95, giving a significant set of 266 probes for which 67 probes had Pr(*S* < 0|𝒟) = 1 (**Fig. 5**). After mapping these probes to genes, a total of 252 genes were identified with the proportion of false positives < 5%. The results of gene ontology (GO) overrepresentation tests showed that these genes were enriched in the molecular function of IgG binding (*P* = 0.032 after Bonferroni correction). Moreover, we detected 45 genes that had Pr(*S* > 0|𝒟) > 0.95 (Fig. 5), which were enriched in the molecular function of α_1_-adrenergic receptor activity (*P* = 0.048) and potassium channel activity (*P* = 0.016). These results are consistent with a previous review^56^ that a proportion of genes showing evidence of selection were significantly enriched in the function of immunity, receptor and potassium channel activity.

**Figure 5:**
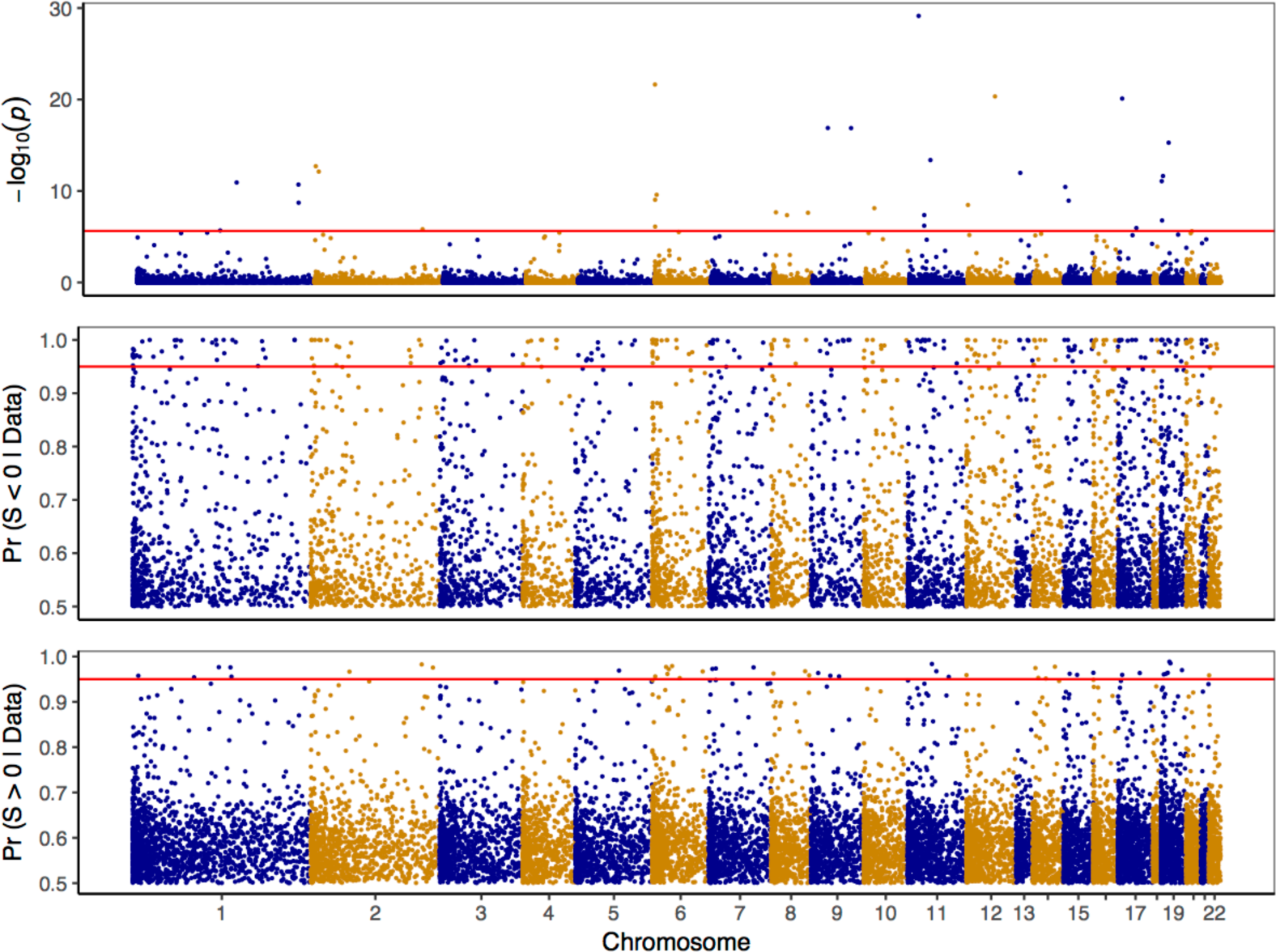
Genome-wide evidence of selection from the p-values to test against *S* = 0 and the posterior probability of *S* < 0 or *S* > 0 for 21,303 probes in the CAGE data after QC. The red line shows the significant threshold of 0.05 after Bonferroni correction (p-value = 2.3×10^−6^) or 0.95 for the posterior probabilities.

### The directions of parameter *S* under different types of natural selection

Besides detecting selection and quantifying its strength on the trait-associated SNPs, the sign of *S* allows us to further infer the type of selection. To demonstrate this, we used forward simulations (**Online Methods**) to simulate common types of natural selection for a quantitative trait by relating the normally distributed phenotype to fitness through a hypothetical function (**Fig. 7**, top row). In the last generation of selection, the relationship between the variance 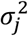 in the effect of coded allele and its frequency showed different patterns across different types of selection (**Fig.7**, bottom row). As expected, when all the variants were selectively neutral, 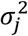 was uniformly distributed across MAF (*S* = 0). Under stabilizing selection,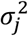 was negatively related to MAF (*S* < 0), a result of purifying trait-associated variants with large effect size which was deleterious to fitness through pleiotropy (also known as negative selection). Both directional (in either direction) and disruptive selection led to a positive relationship between 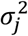 and MAF (*S*> 0). This is because in both cases, alleles with favourable effects increased in frequency due to positive selection, so that high MAF bins were enriched with derived alleles of large effect. The difference is that disruptive selection kept the alleles with large effects at the intermediate frequencies, while directional selection persistently drove them toward fixation, resulting in a sigmoidal or convex shape of the relationship between 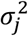 and MAF (**Supplementary Fig. 12**). In conclusion, estimate of *S* is informative to detect the signature of natural selection and is able to distinguish stabilising selection from directional and disruptive selection for a trait. At the level of genetic variants, a negative (positive) value of *S* is indicative of negative (positive) selection on the variants associated with the trait.

## Discussion

We infer the action of natural selection on a complex trait from the signature left in the genetic architecture – the relationship between effect size and MAF. We introduced a method to simultaneously estimate the SNP-based heritability, polygenicity and the relationship between effect size and MAF using all genome-wide SNPs. In contrast to the contemporary methods that use independent SNPs that are significantly associated with traits^3,37,57,58^, our method accounts for genome-wide SNP effects jointly and therefore has more statistical power to detect the signature of selection for polygenic traits. Results of the simulations using real genotypes showed that our estimate of the relationship (*S*) is unbiased when the causal variants are observed; otherwise, the estimate tends to be conservative depending on the LD between SNPs and the causal variants (**Fig. 1**). We detected significant signatures of natural selection (S ≠ 0) for 23 out of 28 complex traits in the UKB data, with the strongest selection signals from the reproductive and cardiovascular trait-associated SNPs, followed by those associated with height, handgrip strength, lung function and other anthropometric traits as well as educational attainment (**Fig. 3**). Our findings are in line with an increasing body of literature supporting the hypothesis of widespread polygenic selection on standing variants in complex traits^4,39,40,59,60^. Together with the high prevalence of selection signals across traits (23/28 = 82%), our observation of high degree of polygenicity (~6% on average) underlines the role of pleiotropy in the action of natural selection.

In the analysis of the UKB data, all the significant estimates of *S* for 23 traits were negative (**Fig. 3**), consistent with a model of negative selection (**Fig. 7**). The evidence of negative selection against the trait-associated variants has been previously reported in some of these traits, such as height and BMI^36^. A recent study on ~110,000 Icelanders also detected negative selection on EA-increasing variants over recent generations, as a result of delayed reproduction and fewer children for the people with higher EA^61^. To support our results on some of the other traits, we used the imputed SNPs based on a reference panel constructed by Haplotype Reference Consortium^62^ to estimate the genetic variance across SNPs that are stratified by MAF and LD scores (GREML-LDMS^36^). We found that the genetic contribution of rare SNPs (MAF < 0.01) is disproportional to that of common SNPs (MAF > 0.01) in height, BMI, WHR, and diastolic blood pressure (**Supplementary Table 6**). These results also suggest that negative selection is pervasive across traits, in line with the conclusion drawn from the BayesS analysis.

In the analyses of CAGE data, we identified 30 genes showing significant signatures, all of which had negative *Ŝ* (**Fig. 6**). With a less stringent criterion for hypothesis testing, we identified additional 267 genes but only 45 of them had positive *Ŝ* (**Fig. 5**). These results again suggest the predominant role of negative selection in the human genome^59,63,64^ and support the hypothesis that gene expression evolves primarily under stabilizing selection^65,66^. The genes that showed evidence of negative selection in our analysis may be functionally important and may deserve a downstream study. There are gene-level metrics, such as Residual Variation Intolerance Score (RVIS)^67^ and Gene Damage Index (GDI)^68^, aiming to prioritize genes for disease involvement based on the functional variation within a gene, which can be used to infer the strength of natural selection on the (coding) sequence of the gene. In contrast, our method interrogates genome-wide SNPs to detect signals of selection on the SNPs associated with the expression level of a gene, which largely depend on the trans-effects for polygenic transcripts. We found that *Ŝ* is not correlated with either RVIS or GDI (**Supplementary Fig. 13**). However, some genes indeed showed strong evidence of selection in both lines. For example, *HERC2* (*Ŝ* = -1.16, RVIS = -5.99) is a pigmentation-related gene which has been suggested a target of recent selection^69,70^. In addition, genes with significantly negative *Ŝ* generally also had low GDI, which is considered to be an indicator of the relative biological indispensability^68^. We do not expect to detect all genes that are known to be under selection, such as the lactase gene *LCT* (*Ŝ* = 0.221, s.e. = 0.317). One possible reason is that the signatures of selection for these genes are concentrated in the cis-regions and therefore might be diluted when we use all genome-wide SNPs to estimate *S*.

**Figure 6:**
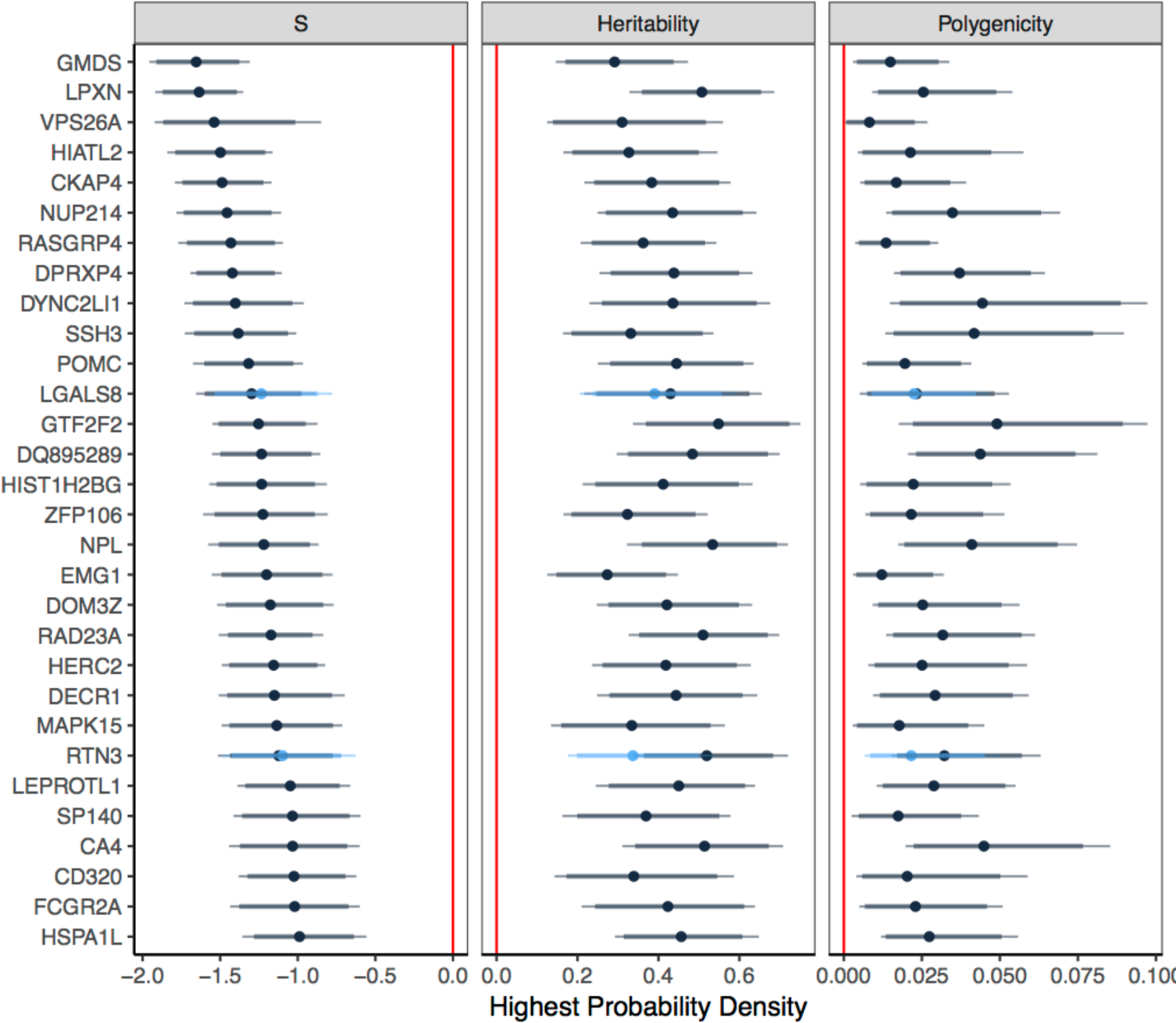
Estimation of the genetic architecture parameters for 30 genes (corresponding to 32 probes) with significant *Ŝ* (p-value < 0.05/21,303) in the analysis of CAGE data. Results are the posterior modes with credible intervals obtained from the nested BayesS model. The bold line represents 95% credible interval (highest posterior density, HPD) and the thin line represents 90% credible interval. Polygenicity is defined as the proportion of 200-Kb windows with nonzero effects in the genome. The light colour shows the results of the second probe tagging one gene.

We conclude with several caveats. First, the polygenicity estimate (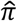) only partially reflects the actual fraction of causal variants since SNPs can possess nonzero effects through LD with unobserved causal variants. Nevertheless, 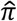 can be used to compare the levels of polygenicity across traits. Second, the power of detecting a signal of natural selection (i.e. testing against *S* = 0) may improve if whole-genome sequence (WGS) or imputed sequence data, which include a large number of rare variants, are used in the analysis. However, it is computationally challenging to run BayesS on all the WGS variants in a large cohort like UKB, a common problem in the analysis of individual-level data with Bayesian methods. Depending on the sparsity of the genetic signals, the nested BayesS provides a possibility to run the analysis in a manageable amount of time but would still require a huge amount of memory to store the genotype matrix for the WGS variants. A more practical approach is to model the genetic architecture using summary statistics. Finally, in the simulation we observed a small inflation in estimating *S* using the nested BayesS model, when all causal variants were genotyped and the true *S* is positive (**Supplementary Fig. 7**). This suggests that the positive *Ŝ* may be slightly overestimated in the CAGE data analysis, but it would not change our conclusions since there was no significant positive *Ŝ*. Given that most complex traits have negative estimates of the relationship between effect size and MAF, we expect to discover more rare variants of large effect in future GWAS using WGS or imputed data.

**Figure 7:**
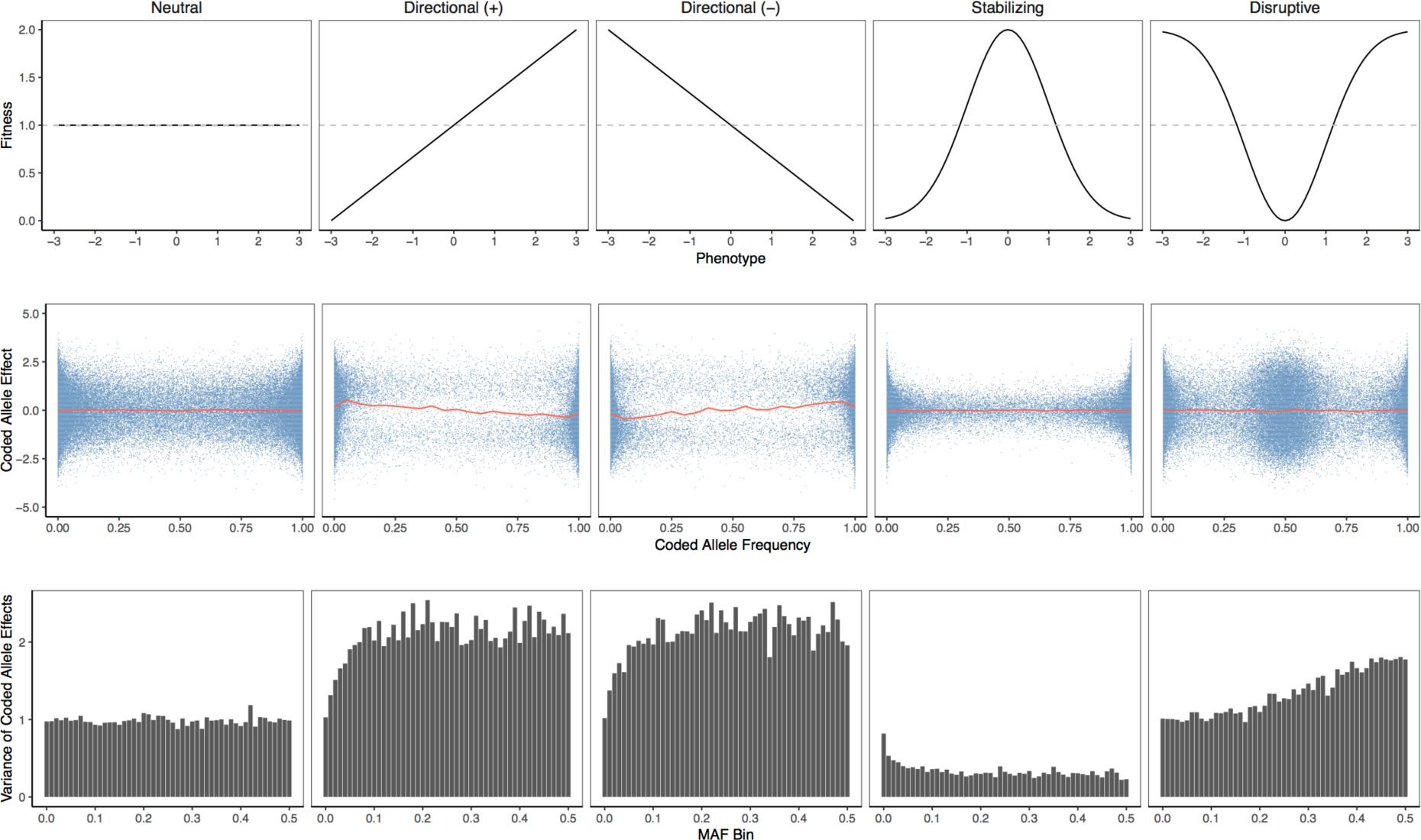
Forward simulations with different types of selection. A quantitative trait was generated based on a simulated chromosome segment of 10Mb (5% causal and 95% neutral mutations in each generation). The trait heritability was 0.1. The top row shows the functions used to relate the phenotype (normally distributed) to fitness in different modes of selection: neutral variation, directional selection with the phenotype positively (+) or negatively (-) correlated to fitness, stabilizing selection and disruptive selection. The 2^rd^ row shows the joint distributions of the coded allele effects and frequencies, where the coded allele at each causal variant was chosen at random from the derived and ancestral alleles, and the red line shows the means of coded allele effects in allele frequency intervals of 0.05. The bottom row shows the relationships between the variance of coded allele effects and MAF. Results were collected from 200 replicates of simulation.

## Online Methods

### The BayesS model

BayesS is a Bayesian multiple regression that simultaneously fits all the SNP effects as random:

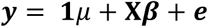

where ***y*** is the vector of phenotypes, *μ* is the fixed effect, **X** is the matrix of SNP genotype scores centred by the column means, ***β*** is the vector of SNP effects, and ***e*** is the residuals. The fixed effect has a flat prior: *μ* ∝ constant. In practice, we fitted principal components and other covariates as well in the model as fixed effects. It is common to standardize the SNP genotypes such that each column of **X** has variance one. But we do not standardize the SNP genotypes, as the standardization implicitly assumes a strong negative relationship between SNP effect size and MAF (*S*= -1)^36,71–73^. As shown in Method Overview, we assume that the SNP effect *β_j_* has a hierarchical mixture prior

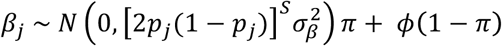

where *ϕ* is a point mass at zero and *π*, the proportion of SNPs with nonzero effects, is the polygenicity. We allow data to dominate the inference on polygenicity by assuming a uniform prior

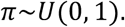

The variance of SNP effects, which quantifies our prior belief on the effect size, is modelled to be related to MAF*p*_*j*_ through *S*, which is assumed to have a normal prior

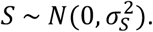

Namely, we *a priori* believe a selectively neutral model with some certainty (quantified by the given variance) to allow the detection of selection to be driven by the data. We set 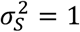 as the prior in the analysis of UK Biobank traits, but a more informative prior 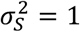 was used in the analysis of CAGE gene expressions to shrink noise heavier toward zero given the much smaller sample size. The prior for the common variance factor is

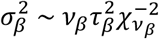

where *v*_*β*_ = 4 and 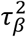 is computed utilizing the characteristic of the distribution: if 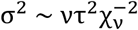, then 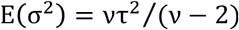. Rearranging the equation gives

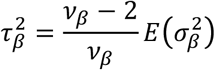

where

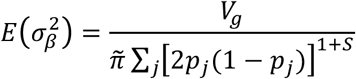

with 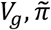 and 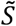 are the prior knowledge of the genetic variance, *π* and *S*. To remove the dependence of the hyperparameter 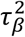 on the prior values of the genetic variance, *π* and _S_, we compute 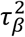 deterministically using the sampled values of these parameters for the first 2,000 MCMC cycles, and then set 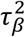 to the average value across these cycles. Likewise, the prior for the residual variance is

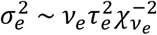

where *v*_*e*_ = 4 and 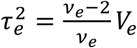 with *V_e_* a prior knowledge of the residual variance. Note that when *S* = 0, our model becomes BayesCπ^19^, a method that has been widely used for genomic prediction in agriculture, or Bayesian Variable Selection Regression (BVSR) in statistics literature^74^. The sampling scheme of the parameters is given in the **Supplementary Note**.

The nested BayesS model is developed based on a previously published method, BayesN^75^, to speed up computation when a large number of SNPs is included in the analysis. In the nested BayesS, the genome is partitioned into *W*-Kb non-overlapping segments. Each window *a priori* has *k* SNPs with nonzero effects, where *W* and *k* are some given numbers. SNPs in the same window are individually modelled as in BayesS as well as collectively considered as a window effect with a normal-zero mixture prior. Remarkable speedups are obtained by “jumping” fast over the windows with zero effect, focusing solely on the windows that harbour genetic signals. Thus, the reduction in computing time is inversely related to the polygenicity, which is defined here as the proportion of segments with nonzero effects. When the causal variants are not observed, choosing *k* > 1 may lead to better performance in parameter estimation than BayesS, as it refines the signal of causal variant by allowing the flanking SNPs to jointly capture its effect. Details on the nested BayesS and the comparison with the standard BayesS are given in the **Supplementary Note**.

### Estimation of heritability

We estimate the SNP-based heritability using the sampled values of SNP effects in MCMC. By definition, the genetic variance is the variance of the genetic values across individuals. In each MCMC cycle, we calculate the genetic values for each individual (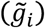) using SNPs with sampled nonzero effects 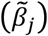:

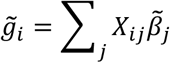

Then, the genetic variance in the current cycle is

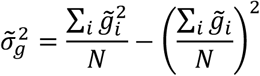

Conditional on the sampled value of residual variance (σ|), the SNP-based heritability is

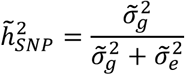

The mean over all cycles after burn-in is the estimate of heritability

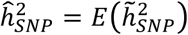

The standard deviation of the MCMC samples gives the standard error of the estimate and the highest probability density gives the credible interval for posterior inference.

### ARIC simulation analysis

The simulation based on Atherosclerosis Risk in Communities (ARIC) cohort^33^ was used for testing the methods. We used PLINK 1.9^76^ to carry out standard quality control (QC) procedures on the dataset, including removal of SNPs with missingness > 5%, Hardy-Weinberg equilibrium test *P <* 10^−6^, or MAF < 1%, and removal of individuals with missing genotypes < 1% and genetic relationship < 0.05 estimated from all SNPs after QC using GCTA-GRM^77^. After all the QC steps, a total of 12,942 unrelated individuals and 564,959 SNPs remained. A quantitative trait was simulated by choosing 1,000 SNPs at random as causal variants with their effects sampled from a standard normal distribution. To simulate a spectrum of relationships between MAF and effect size, the effect size was multiplied by 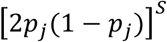where *S* = −2,−1,0,1, or 2, representing negative to positive relationship between effect size and MAF including the case of independence when *S* = 0. An individual phenotype specific to a given value of *S* was generated by adding a random normal residual with the variance identical to the genetic variance, giving each simulated trait a heritability of 0.5. The simulation process was repeated for 100 times. BayesS and the nested BayesS were used to analyse the simulated data with and without the causal variants in the model. To evaluate the robustness of our method to the starting values of parameters, we used an arbitrary value of 0 for *S*, 0.2 for heritability, and 0.05 for *π*, respectively to start the MCMC. In the nested BayesS, the length of window was set to be 200-Kb with 2 SNPs *a priori* fitted in the model. It is noteworthy that the distribution of genetic variance explained by each causal variant was not identical for different scenarios of *S* in the true model. Under HWE, the genetic variance at locus *j* is 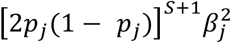 with *β_j_* ~ *N*(0,1) in the simulation. Compared to a trait with *S* < 0, a trait with *S* > 0 has a larger proportion of loci each explaining a small proportion of variance, given an identical distribution of MAF at the causal variants between traits (**Supplementary Fig. 14**). Thus, in the scenario of *S* > 0, it is more difficult to capture the causal variants by SNP markers if causal variants are not observed.

### Analyis of the UK Biobank data

We have access to 46 complex traits in the UK Biobank^26^, where the phenotype data were collected from over 500,000 individuals aged between 40 and 69 across the United Kingdom. The interim release contains genotypes for 152,736 samples at 806,466 SNPs on a customized Affymetrix Axiom array after QC procedures^78^. We selected a subset of 140,408 individuals that had a self-reported gender identical to the genetically inferred gender and a European ethnicity derived from a principal component analysis together with self-reported ethnicity. Furthermore, we removed individuals with genomic relatedness > 0.05 estimated from all SNPs using GCTA-GRM^77^ and SNPs with genotype missing rate > 5%, Hardy-Weinberg equilibrium test *P*< 10^−6^, or MAF < 1%. The final data set consisted of 126,752 individuals of European ancestry with 483,634 common SNPs (MAF > 1%). After removal of 5 duplicated traits and 5 traits with sample size *(N)* < 20,000, we had 36 traits remained for analysis, including 32 quantitative traits (anthropometric, cardiovascular and reproductive), 2 categorical traits - male pattern baldness (MPB) and years of schooling (educational attainment) and 2 diseases - type 2 diabetes (T2D) and major depressive disease (MDD). The sample sizes of the traits are shown in **Supplementary Table 2**, where most traits had *N* > 100,000. The prevalence of T2D and MDD in the sample was 5.35% and 6.70%, respectively. The phenotypes of quantitative traits were standardized within each sex group after regressing out the age effect. For educational attainment, the years of schooling are preadjusted by sex, a third order polynomial of year-of-birth and year-of-birth by sex interactions. We used BayesS for the analysis, where the first 20 principal components (PC) of GRM were fitted as fixed effects to account for the effects due to population stratification. For the disease traits, sex and age were fitted as covariates in addition to PCs, and for MPB, only age was fitted as the additional covariate.

### Consortium for the Architecture of Gene Expression (CAGE) data set

We analyzed the mRNA levels for 36,778 transcript expression traits (probes) from the Consortium for the Architecture of Gene Expression (CAGE)^27^ data set using the nested BayesS method. The CAGE data comprised of measurements from 36,778 gene expression probes in peripheral blood, with a subset of 1,748 unrelated (genomic relatedness > 0.05) European individuals from the total 2,765 individuals used for this analysis. Full details of the cohorts contributing to CAGE, and their sample preparation, normalization and genotype imputation are outlined in Lloyd-Jones *et al.*^27^. Briefly, the quantification of gene expression for each cohort was performed using the Illumina Whole-Genome Expression BeadChips. The gene expression levels in each cohort were initially normalized, followed by a quantile adjustment to standardize the distribution of expression levels across samples. We corrected for age, gender, cell counts and batch effects as well as hidden heterogeneous sources of variability. The rank-based inverse-normal transformation was used to normalize the measurements for each probed to be normally-distributed with mean 0 and variance 1. Probes measuring expression levels of genes located on chromosomes X and Y were removed from the analysis. The initial CAGE dataset consisted of seven unique cohorts that were genotyped on different SNP arrays. Therefore, genotype data were imputed to the 1000 Genomes Phase 1 Version 3 reference panel^79^, resulting in 7,763,174 SNPs passing quality control of which 1,066,738 SNPs overlapped with HapMap3 and were used for analysis.

### Forward simulation for different types of natural selection

We ran forward simulations using SLiM^80^ to confirm that the relationship between effect size and MAF is subject to different types of natural selection. We simulated a 10-Mb region where new mutations occurred with probability of 0.95 to be neutral and of 0.05 to be a causal variant with an effect sampled from a standard normal distribution. The mutation rate was set to be 1.65×10^−8^^81^. The phenotype of an individual was simulated based on the genotypic values at all segregating causal variants in the current generation with a heritability of 0.1. We simulated the evolution of a population of 1,000 individuals over 10,000 generations (this is equivalent to 100,000 generations in a population of 10,000 individuals^82^). The first 5,000 generations were used a burn-in period, where the phenotype did not affect fitness and all variants (including the causal variants) were under neutral variation. From generation 5,001, we related the standardized phenotype with mean zero and variance one to fitness through a hypothetical function that represents different types of selection (**Fig. 7**, top row). For directional selection, the phenotype was positively or negatively correlated to fitness through a simple linear function. For stabilizing selection, we used a normal curve to model that fitness achieved optimum at intermediate phenotype value. For disruptive selection, a reversed normal curve was used to model that the phenotypes at the two tails produced highest fitness. In the last generation of selection, we investigated the joint distribution of effects and frequencies of the derived alleles, the joint distribution of effects and frequencies of the coded alleles (arbitrarily chosen as in reality where derived alleles are unknown), and the relationship between the variance of the coded-allele effects and MAF. We collected results from 200 simulation replicates.

## Acknowledgments

This research was supported by the Australian Research Council (DP160101343), the Australian National Health and Medical Research Council (1107258, 1078901, 1078037, and 1113400), and the Sylvia & Charles Viertel Charitable Foundation (Senior Medical Research Fellowship). R.d.V. acknowledges funding from an ERC consolidator grant (647648 EdGe, awarded to Philipp Koellinger). This study makes use of data from dbGaP (accession number: phs000090.v3.p1) and UK Biobank Resource (Application number: 12514). A full list of acknowledgements to these data sets can be found in the **Supplementary Note**. At the end, we wish to acknowledge The University of Queensland's Research Computing Centre (RCC) for its support in this research.

## Author contributions

J.Y., P.M.V. and R.d.V. conceived the study. J.Y., J.Z. and P.M.V. designed the experiment. J.Z. derived the analytical methods, conducted all analyses and developed the software with assistance and guidance from J.Y., Y.W., M.R.R., L.L.J., L.Y.D., C.Y. A.X. and J.S. L.L.J., A.F.M., J.E.P., G.W.M., A.M., T.E., G.G. and P.M.V. provided the CAGE data. J.Z. and J.Y. wrote the manuscript with the participation of all authors. All authors reviewed and approved the final manuscript.

